# Emergence of wheat blast in Bangladesh was caused by a South American lineage of *Magnaporthe oryzae*

**DOI:** 10.1101/059832

**Authors:** M. Tofazzal Islam, Daniel Croll, Pierre Gladieux, Darren M. Soanes, Antoine Persoons, Pallab Bhattacharjee, Shaid Hossain, Dipali Rani Gupta, Md. Mahbubur Rahman, M. Golam Mahboob, Nicola Cook, Moin U. Salam, Vanessa Bueno Sancho, João Leodato Nunes Maciel, Antonio Nhani Júnior, Vanina Lilián Castroagudín, Juliana T. de Assis Reges, Paulo Cezar Ceresini, Sebastien Ravel, Ronny Kellner, Elisabeth Fournier, Didier Tharreau, Marc-Henri Lebrun, Bruce A. McDonald, Timothy Stitt, Daniel Swan, Nicholas J. Talbot, Diane G.O. Saunders, Joe Win, Sophien Kamoun

## Abstract

In February 2016, a new fungal disease was spotted in wheat fields across eight districts in Bangladesh. The epidemic spread to an estimated 15,741 hectares, about 16% of cultivated wheat area in Bangladesh, with yield losses reaching up to 100%. Within weeks of the onset of the epidemic, we performed transcriptome sequencing of symptomatic leaf samples collected directly from Bangladeshi fields. Population genomics analyses revealed that the outbreak was caused by a wheatLinfecting South American lineage of the blast fungus *Magnaporthe oryzae*. We show that genomic surveillance can be rapidly applied to monitor plant disease outbreaks and provide valuable information regarding the identity and origin of the infectious agent.

---

Outbreaks caused by fungal diseases have increased in frequency and are a recurrent threat to global food security (Fisher et al., 2012). One example is blast, a fungal disease of rice, wheat and other grasses, that can destroy enough food supply to sustain millions of people (Fisher et al., 2012; Pennisi, 2010; Liu et al., 2014). Until the 1980s, the blast disease was not known to affect wheat, a main staple crop critical to ensuring global food security. In 1985, the disease was first reported on wheat (*Triticum aestivum* L.) in Paraná State, Brazil (Igarashi et al. 1986). It has since spread throughout many of the important wheat-producing areas of Brazil and to neighboring South American countries including Bolivia and Paraguay. In South America, blast is now a major threat to wheat production (Goulart et al., 1992; Goulart et al., 2007; Kohli et al., 2011). Currently, wheat blast affects as much as 3 million hectares seriously limiting the potential for wheat production in the vast grasslands region of South America.

Blast diseases of grasses are caused by fungal species from the *Pyriculariaceae* (Klaubauf et al., 2014) and can occur on 50 grass species (Ou, 1985). However, a high degree of host-specificity exists among and within these fungal species (Kato et al. 2000; Klaubauf et al., 2014). In South America, wheat blast is caused by isolates of *Magnaporthe oryzae* (syn. *Pyricularia oryzae*) known as pathotype *Triticum* (Urashima et al., 1993; Kato et al., 2000; Tosa et al, 2006). The rice-infecting isolates of *M. oryzae* are genetically distinct from wheat-infecting isolates and generally do not infect wheat (Prabhu et al., 1992; Urashima et al., 1993; Urashima et al., 1999; Farman 2002; Faivre-Rampant et al. 2008; Tufan et al., 2009; Maciel et al. 2014; Chiapello et al., 2015; Yoshida et al., 2016). Typical symptoms of wheat blast on spikes are premature bleaching of spikelets and entire heads (Igarashi 1990; Urashima 2010). Severely infected wheat heads can be killed, resulting in severe yield losses (Igarashi 1990; Urashima 2010). The disease is generally spread by infected seeds and airborne spores, and the fungus can survive in infected crop residues and seeds (Urashima et al., 1999). Little is known about the physiology and genetics of the wheat blast pathogen, and our understanding of molecular interactions of this pathogen with wheat remains limited.

In February 2016, wheat blast was detected for the first time in Asia with reports of a severe outbreak in Bangladesh relayed through local authorities and the media. Although wheat is not a traditional crop in Bangladesh, its cultivation has expanded in recent years making it the second major food source after rice (Hossain and Teixeira da Silva, 2012). The outbreak is particularly worrisome because wheat blast could spread further to major wheat producing areas in neighboring South Asian countries, thus threatening food security across the region. Here, we report our immediate response to this plant disease outbreak. To rapidly determine the precise identity and likely origin of the outbreak pathogen, we applied field pathogenomics, in which we performed transcriptome sequencing of symptomatic and asymptomatic leaf samples collected from infected wheat fields in Bangladesh (Hubbard et al., 2015; Derevnina and Michelmore, 2015). To promote the project and recruit experts, we immediately released all raw sequence data through a dedicated website OpenWheatBlast (http://www.wheatblast.net). Phylogenomic and population genomic analyses revealed that the Bangladesh wheat blast outbreak was probably caused by isolates belonging to the South American wheat-infecting lineage of *M.( oryzae*. We conclude that the wheat blast pathogen was most likely introduced into Asia from South America.

## RESULTS AND DISCUSSION

### Geographical distribution of the wheat blast outbreak in Bangladesh

The total area of wheat cultivation in Bangladesh in 2016 was about 498,000 ha (Department of Agricultural Extension, Bangladesh). Wheat blast was observed in eight south-western districts viz. Pabna, Kushtia, Meherpur, Chuadanga, Jhenaidah, Jessore, Barisal and Bhola (Fig. 1). Out of a total 101,660 ha of cultivated wheat in those 8 districts, an estimated 15,471 ha (16%) was affected by wheat blast.

**Figure 1.**
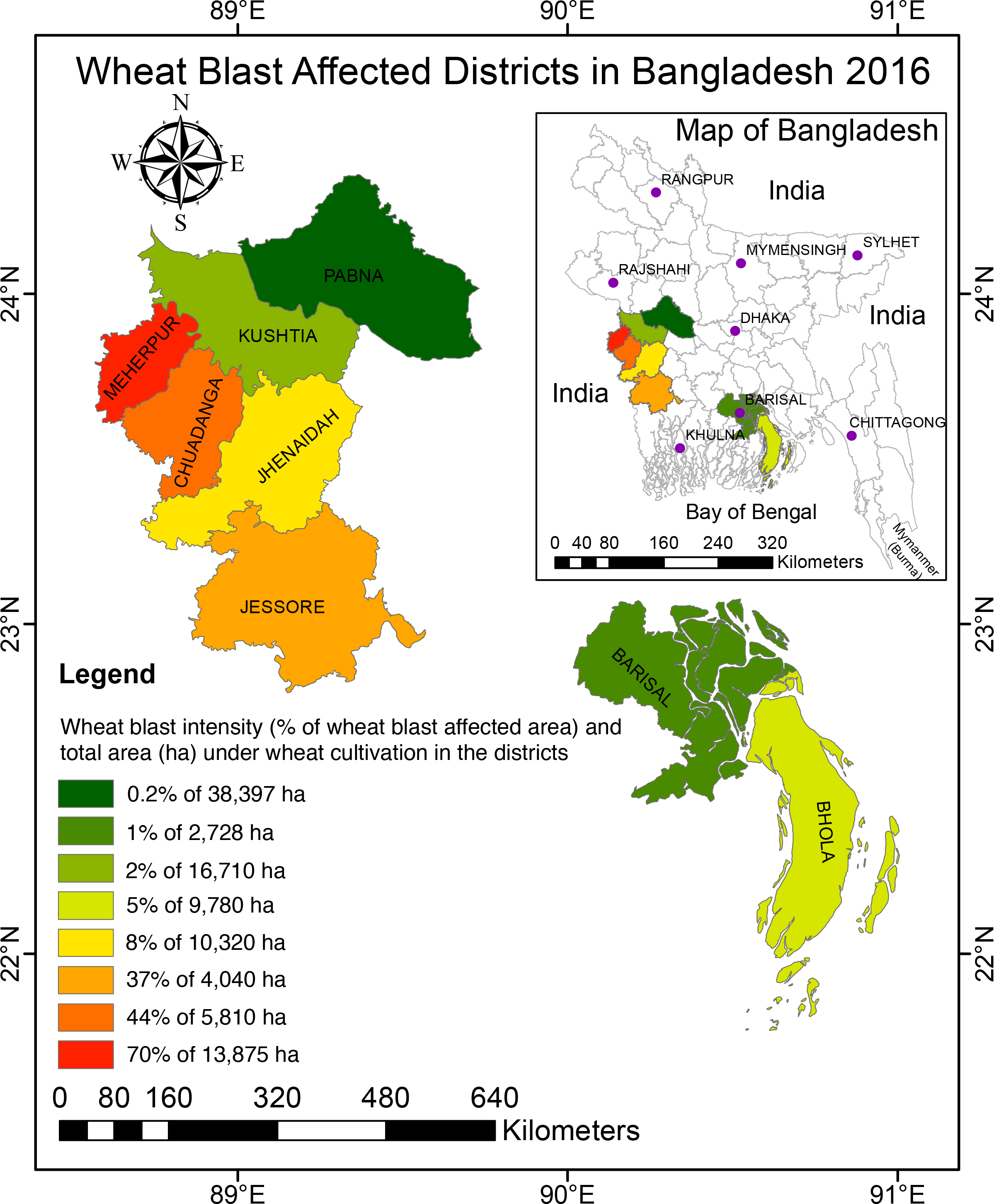
Geographical distribution and severity of the wheat blast outbreak in eight southwestern districts of Bangladesh. The map depicts the intensity of the 2016 wheat blast outbreak across Bangladesh. The % of affected area and the total area (hectares) under cultivation is shown for each district based on the color chart.

The severity of wheat blast and associated yield losses varied among districts. The highest percentage of infected wheat fields was observed in Meherpur (70%) followed by Chuadanga (44%), Jessore (37%), Jhenaidah (8%), Bhola (5%), Kushtia (2%), Barisal (1%) and Pabna (0.2%) (Fig. 1). Yield losses in different affected districts varied. The highest average yield loss was recorded in Jhenaidah (51%) followed by Chuadanga (36%), Meherpur (30%), Jessore (25%), Barisal (21%), Pabna (18%), Kushtia (10%), and Bhola (5%). Although the average yield losses was lower than 51% across districts, yield losses in individual fields were as high as 100%. Importantly, 100% of government-owned Bangladesh Agricultural Development Corporation (BADC) seed multiplication farms in the affected districts (ca. 355 ha) were completely cleared by burning to destroy pathogen inocula by decision of the Ministry of Agriculture (see https://www.youtube.com/watch?v=EmL5YM0kIok). Farmer wheat fields that were severely affected (~10%) were also burned.

### Wheat blast symptoms in the field

To examine disease symptoms in affected wheat fields, we collected samples from the affected districts. Major symptoms associated with the epidemic included completely or partially bleached (dead) spikes similar to symptoms reported for Brazilian wheat blast epidemics (Igarashi et al., 1990; Urashima, 2010). The pathogen attacked the base or upper part of the rachis, severely affecting spikelet formation above the point of infection. Complete or partial bleaching of the spike above the point of infection with either no grain or shriveled grain was common in all areas affected by wheat blast (Fig. 2A–Fig 2C). We commonly observed bleached heads with traces of gray, indicative of fungal sporulation at the point of infection (arrows in Fig. 2A–Fig 2C and Fig. 2G). In severely infected fields, we also found typical eye-shaped necrotic disease lesions with gray centers in leaves of some wheat plants (Fig. 2D) (Igarashi, 1990; Cruz et al., 2015). Head infections during the flowering stage resulted in no grain production (Fig. 2G), whereas infection at the grain filling stage resulted in small, shriveled, light in weight, and discolored (pale) grains (Fig. 2E–Fig. 2F) (Urashima, 2010).

**Figure 2.**
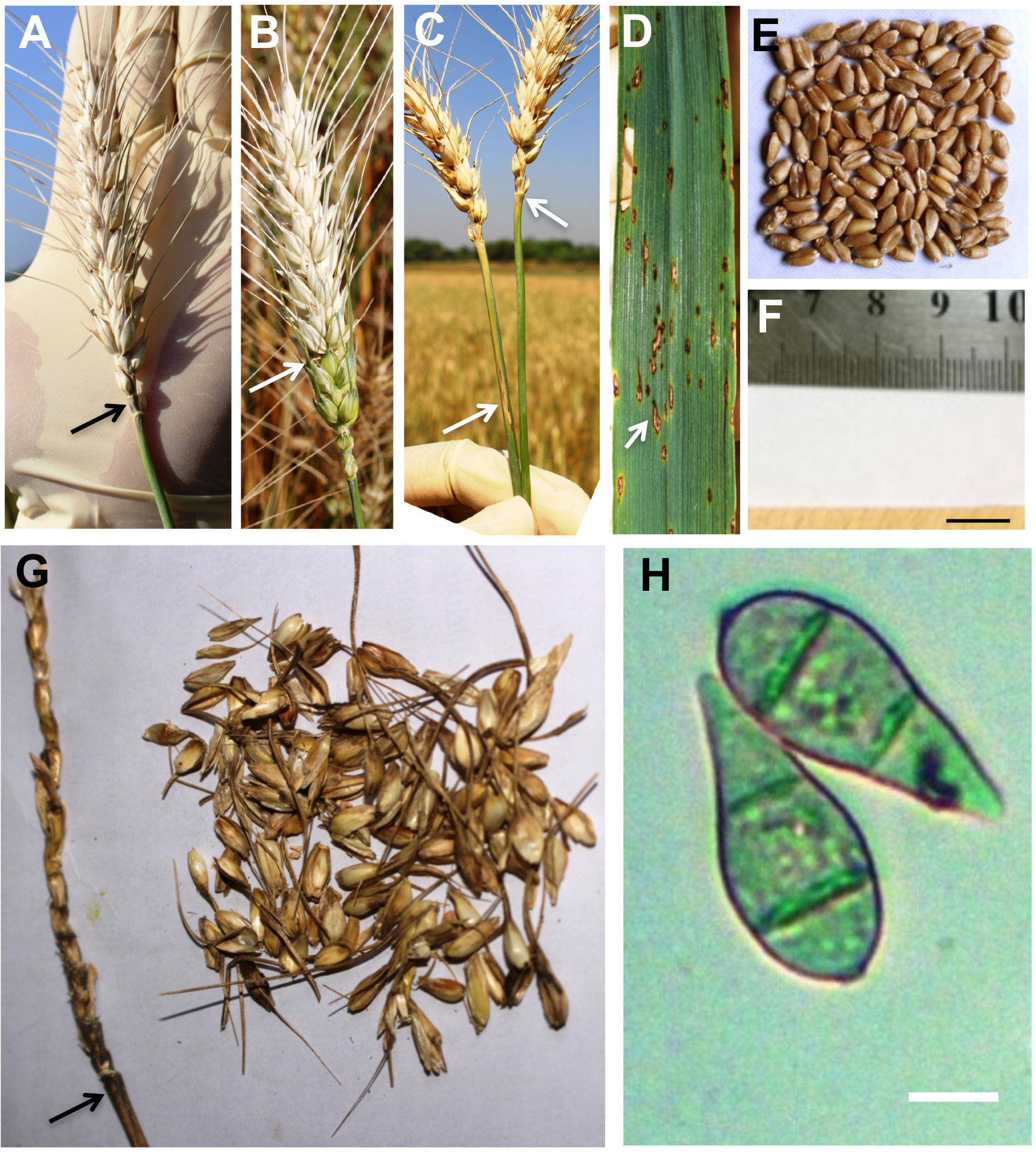
Symptoms of blast disease in wheat and a micrograph of conidia of *Magnaporthe oryzae* isolated from a farmer’s field in Jhenaidah, Bangladesh. Symptoms of blast disease in spikes, leaves, and seeds of wheat in a farmer’s field in Jhenaidah in Bangladesh, and a micrograph showing two conidia of *Magnaporthe oryzae*. (A) A completely bleached wheat spike with traces of gray from blast sporulation at the neck (arrow) of the spike. (B) Complete bleaching of a wheat spike above the point (arrow) of infection. (C) Two completely bleached spikes with traces of gray (upper arrow) and a lesion (lower arrow) from blast sporulation at the base. (D) Typical eye-shaped lesion (arrow) and dark gray spots on a severely diseased wheat leaf. (E) Mild blast disease-affected slightly shriveled wheat seeds. (F) Severe blast-affected shrivelled and pale-colored wheat seeds. (G) A severely infected rachis with dark gray blast sporulation at the neck (arrow) and severely damaged spikelets. (H) Micrograph of two conidia isolated from the infected spike of wheat. Scale bars in E and F = 1 cm and H = 10 mm.

To determine whether the spike and leaf symptoms on wheat were associated with infection by blast fungi (*Pyricularia* and related genera from the *Pyriculariaceae* sensu Klaubauf et al., 2014), we examined infected plant samples using a light microscope. A hallmark of blast fungi is the production of asexual spores that have a specific morphology consisting of three-celled pyriform conidia (Klaubauf et al., 2014). Microscopic analyses revealed that gray colored lesions observed on both spikes and leaves carried large numbers of three-celled pyriform conidia (Fig. 2H). This indicates that the fungus present in these lesions belongs to the *Pyriculariaceae*. Molecular taxonomy tools are however needed to determine the species identity.

### Transcriptome sequencing of wheat leaf samples from Bangladeshi fields

We used field pathogenomics (Hubbard et al., 2015) to identify which blast fungus species was present in infected wheat fields in Bangladesh. We collected samples of both symptomatic and asymptomatic leaves from wheat fields in different regions of Bangladesh, including Meherpur and Jhinaidaha districts, and extracted total RNA from four pairs of symptomatic (samples 2, 5, 7 and 12) and asymptomatic samples (samples F2, F5, F7 and F12) (Table S1). We prepared and sequenced RNA-seq libraries using Illumina technology, yielding 68.8 to 125.8 million 101 bp pair-end reads with an average insert size of 419 bp. Next, following data trimming, we aligned high-quality reads to both the *M. oryzae* wheat blast fungus BR32 and wheat genomes (International Wheat Genome Sequencing Consortium, 2014; Chiapello et al., 2015). Sequence reads from all samples with disease symptoms aligned to the BR32 genome, ranging from 0.5% to 18.6% of the total reads (Fig. 3A). By contrast, only a minor proportion of the reads from the asymptomatic samples aligned to the BR32 genome (range: 0.003% to 0.037%, Fig. 3A–Fig. 3C). Between 37.7% and 86.5% of total reads aligned to the wheat genome sequence (Fig. 3A). These analyses revealed that *M. oryzae* is present in symptomatic (infected) wheat samples from Bangladesh.

**Figure 3.**
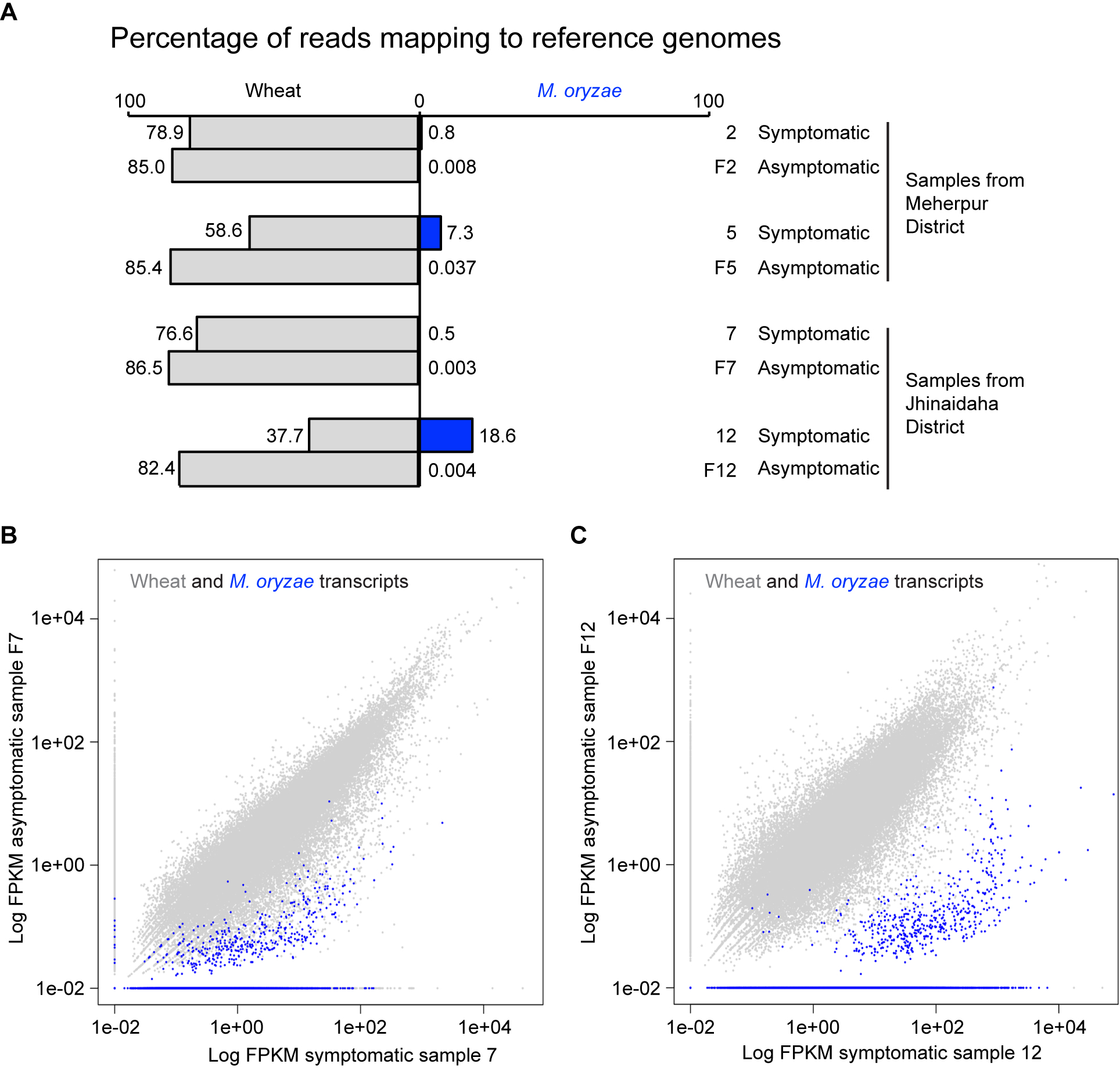
Transcriptome sequencing of infected leaves from farmer fields reveals *Magnaporthe oryzae* transcripts in symptomatic samples. (A) Comparison of sequence read mapping data from the four sample pairs to the genomes of wheat blast fungus *M. oryzae* BR32 (in blue) and wheat (light gray). (B and C) Scatter plot of Fragments Per Kilobase of transcript per Million (FPKM) values from sample pair 7-F7 (B) and 12-F12 (C) aligned to the combined transcriptomes of wheat and *M. oryzae* BR32. Transcripts from wheat (100,344) are shown in light gray and transcripts from *M. oryzae* BR32 (14,349) are shown in blue.

### The Bangladesh wheat outbreak was caused by a wheatLinfecting South American lineage of *M. oryzae*

We used phylogenomic approaches to determine how related the fungal pathogen detected in wheat leaf samples from Bangladesh is to *M. oryzae* lineages infecting cereals and grasses. We also performed population genomics analyses to gain insight into the geographic origin of these Bangladeshi isolates using a set of sequences from wheat-infecting *M. oryzae* isolates collected in Brazil over the last 25 years. We first determined the taxonomic affiliation and phylogenetic position of wheat-infecting Bangladeshi samples. To this aim, we extracted predicted transcript sequences from the assembled genomic sequences of 20 *M. oryzae* strains isolated from infected rice (*Oryza sativa*), wheat (*Triticum aestivum*), foxtail millet (*Setaria* spp.), *Eleusine* spp., *Lolium* spp., and *Eragrostis* spp. (Chiapello et al. 2015; this study; see Table S1 for full details). We identified 2,193 groups of sequences with orthologous relationships across the 20 reference transcriptomes and the two Bangladeshi isolates that had the largest number of genes represented in their transcriptomic sequences. We aligned orthologous transcripts, processed alignments and inferred a maximum likelihood genealogy based on the concatenated sequences using RAxML (Stamatakis, 2014). The Bangladeshi isolates clustered with high bootstrap support (>90%) with wheat-infecting isolates of *M. oryzae* (Fig. 4A), indicating that the emergence of wheat blast in Bangladesh was caused by isolates belonging to the known *M. oryzae* wheat-infecting lineage, and not by an unknown *Pyriculariaceae* species nor a novel *M. oryzae* lineage.

**Figure 4.**
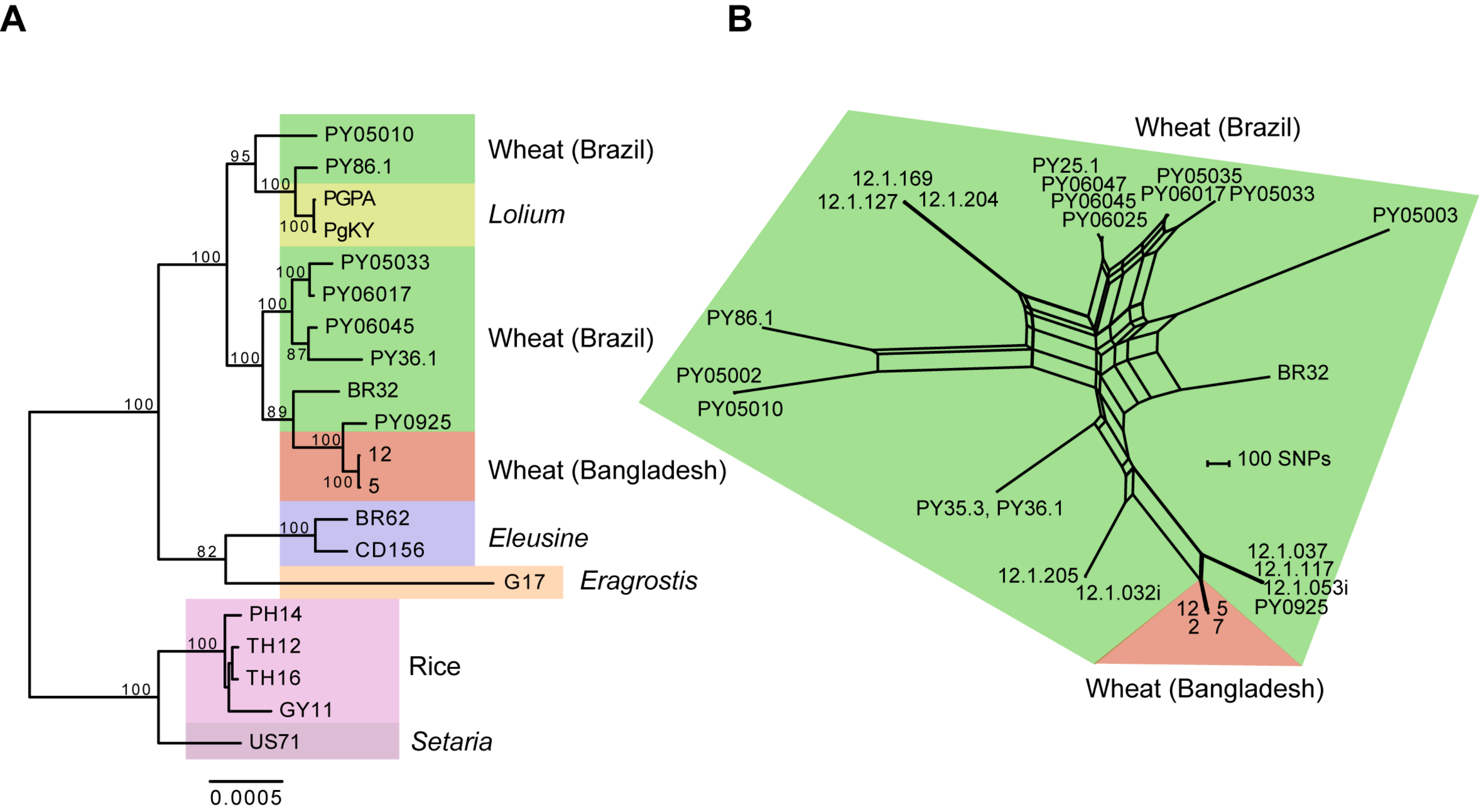
The origin of the Bangladesh wheat blast fungus. (A) Maximum likelihood genealogy inferred from the concatenation of aligned genomic data at 2,193 orthologous groups of predicted transcript sequences. Scale bar represents the mean number of nucleotide substitutions per site. (B) Population genomic analyses of transcriptomic single nucleotide polymorphisms among *M. oryzae* isolates from wheat in Brazil and Bangladesh. The network was constructed using the Neighbor-net algorithm. The scale shows the number of informative sites.

Given that the Bangladesh outbreak was caused by isolates related to known wheat-infecting lineages of *M. oryzae*, our next step was to infer the most likely migration route of the disease using population genomics. We performed population genomics analyses using transcriptomic single nucleotide polymorphisms (SNPs) identified by aligning sequence reads to the *M. oryzae* reference genome 70-15 (Dean et al., 2005). We included all four symptomatic samples from Bangladesh and a diverse collection of 23 *M. oryzae* wheat-infecting isolates sampled from Brazil, the main wheat growing country affected by wheat blast (Table S1). As the wheat blast isolates from Brazil were sequenced from genomic DNA, we restricted the analyses to transcriptomic SNPs genotyped at high confidence in the symptomatic Bangladesh sample 12, retaining a total of 15,871 SNPs. Since the reproductive mode of wheat blast populations can be mixed, including both sexual and asexual reproduction (Maciel et al., 2014), we chose to build a Neighbor-net network that takes into account potential recombination among genotypes. The network analyses identified small groups of near-clonal genotypes (e.g. isolates 12.1.205 and 12.1.032i) whereas all other isolates appeared genetically distinct and displayed reticulate evolution. The Bangladesh outbreak isolates grouped as a near-clonal genotype that was most closely related to a group of Brazilian wheat-infecting isolates from Minas Gerais, Sao Paulo, Brasilia and Goias (strains PY0925, 12.1.053i, 12.1.117, 12.1.037, respectively).

Thus, our rapid open source genomic surveillance approach has revealed the precise identity of the infectious Bangladeshi fungus as the known wheat-infecting *M. oryzae* lineage and indicated it most likely originated from South America. This finding calls for intensive monitoring and surveillance of the wheat blast pathogen to limit its further spread outside South America and Bangladesh. In addition, our finding indicates that the knowledge acquired to manage wheat blast in Brazil using disease resistant cultivars (Anh et al., 2015; Cruz, et al., 2016; Ha et al., 2016) and fungicides (Pagani et al., 2014; Castroagudin et al., 2015) can be directly applied to the Bangladeshi epidemic.

## MATERIALS AND METHODS Field data

The date of first incidence of disease and areas of wheat cultivation and blast-infected fields in different districts of Bangladesh were obtained from the Department of Agricultural Extension (DAE) of Bangladesh. A second data set on the first incidence of disease, the cultivated wheat variety, date of sowing, sources of seeds, and yield loss were directly collected from the farmers (n = 100) of the most severely infected wheat blast district, Meherpur, through face-to-face interviews of randomly selected farmers after harvesting the crop. Among 15,471 ha with wheat blast in Bangladesh, the disease incidence in the Meherpur district alone had 9,640 ha, approximately 62% of the total wheat blast area in the country.

## Transcriptome sequencing of field collected samples

Leaf blades from wheat displaying blast symptoms and those with no symptoms were harvested from the same fields, cut into thin strips (approx. 0.5 × 1.0 cm), and immediately stored in 1 ml RNAlater solution (Thermofisher Scientific, Basinstoke, UK). Total RNA was extracted from the samples using the RNeasy Plant Mini kit (Qiagen, Manchester, UK) following the manufacturer’s instructions. The amount and the quality of RNA samples were determined using the Agilent 2100 Bioanalyzer (Agilent Technologies, Edinburgh, UK). cDNA libraries were prepared using the Illumina TruSeq RNA Sample preparation Kit (Illumina, Cambridge, UK). Library quality was confirmed before sequencing using the Agilent 2100 Bioanalyzer (Agilent Technologies, Edinburgh, UK). The libraries were sequenced on the Illumina HiSeq 2500 system (Illumina) operated by The Genome Analysis Centre, UK, producing 101-bp paired-end reads. The reads were mapped to the genomes of wheat and wheat blast strain *M. oryzae* BR32 using the TopHat software, version 2.0.11 (Kim et al., 2013), and Fragments Per Kilobase of transcript per Million (FPKM) values of mapped reads to the transcriptomes were calculated using Cufflinks, version 2.1.1 (Trapnell et al., 2012). De novo assembly of transcriptomes were performed using sequence reads from each sample with Trinity software, version 2.06 (Grabherr et al., 2011). Within days of sequencing, the data was made public on Open Wheat Blast (http://www.wheatblast.net). A timeline from sample collection to population and phylogenomic analysis is provided on the open wheat blast website (http://s620715531.websitehome.co.uk/owb/?p=485).

## Population and phylogenomic analyses

We used predicted transcript sequences extracted from the assembled genomic sequences of 20 *M. oryzae* isolates collected on infected leaves of rice (*Oryza sativa*), wheat (*Triticum aestivum*), foxtail millets (*Setaria* spp.), *Eleusine* spp., *Lolium* spp., and *Eragrostis* spp. (Table S1; Dobinson et al., 1993; Kang et al., 1995; Tosa et al., 2007; Maciel et al., 2014; Chiapello et al., 2015). We used ProteinOrtho (Lechner et al., 2011) to identify groups of sequences with orthologous relationships across the 20 reference transcriptomes and each of the Bangladeshi transcriptomes. We identified 983, 3,250, 501 and 3,413 groups of orthologous sequences across the reference transcriptomes from samples 2, 5, 7, 12, respectively. Only the two Bangladeshi isolates that had the largest number of orthologous sequences were retained for further analysis (samples 5 and 12). The consensus set of orthologous transcripts across the 22 transcriptomes included 2,193 groups of sequences. We aligned orthologous groups of sequences using MACSE, with default parameters (Ranwez et al., 2011).

We removed codons with missing data or alignment gaps. We excluded transcript alignments for which >0.5% sites corresponded to singletons or doubletons exclusive to the Bangladeshi isolates, suggesting erroneous assignment of predicted sequences to *M. oryzae* BR32 transcripts or sequencing errors in transcript assemblies. We also excluded the regions corresponding to the first 30 and last 16 codons and treated ambiguities as missing data. Maximum likelihood phylogenetic inference was performed on the concatenated sequence of 1,923 orthologs (2,676,792 bp in total), using the GTRGAMMA model in RAxML v. 8.1.17 with 100 bootstrap replicates (Stamatakis, 2014). The maximum likelihood genealogy was mid-point rooted along the longest branch, which was the branch connecting the foxtail millet-and rice-infecting lineages to other lineages.

For population genomic analyses, we identified transcriptomic SNPs based on short read alignments against the *M. oryzae* reference genome 70-15. We mapped quality-trimmed Illumina short read data generated from RNA using TopHat v. 2.0.14 (Trapnell et al., 2012). For all completely sequenced genomes, we aligned quality-trimmed Illumina short read data against the reference genome 70-15 using Bowtie v. 2.2.6 (Langmead and Salzberg, 2012). For all strains collected from the Bangladesh outbreak, transcriptomic sequences were aligned using TopHat v. 2.0.14. We identified variants in the genomes of the different strains using the Genome Analysis Toolkit (GATK) version 3.5 from the Broad Institute (DePristo et al., 2011). We used a two-step variant-calling according to best practice guidelines of GATK. We first called raw variants with local reassembly of read data using HaplotypeCaller. All raw variant calls were jointly genotyped using GenotypeGVCFs. We used SelectVariants to subset the variant calls to contain only SNPs. Then, SNPs were hard-filtered using the following criteria: QUAL ≥ 5000.0, QD ≥ 5.0, MQ ≥ 20.0, −2.0 ≤ ReadPosRankSum ≤ 2.0, −2.0 ≤ MQRankSum-upper ≤ 2.0, −2.0 ≤ BaseQRankSum ≤ 2.0. Furthermore, we only retained SNPs genotyped in at least 90% of all strains and genotyped the Bangladesh sample 12 (Table S1). We used SplitsTree v. 4.14.2 to generate a NeighborNet network from Brazilian and Bangladesh wheat blast strains (Huson and Bryant, 2006). To build the network, we used uncorrected *p* distances calculated from the SNP supermatrix. The network was drawn based on Equal Angle splits.

## ACKNOWLEDGEMENTS

We are thankful to Director General of the Department of Agricultural Extension, Md. Shahidul Islam of Unnnayan Dhara in Jhenaidah, Md. Aminullah of BADC Nurnagar Farm in Chuadanga, and Md. Shabab Mehebub, East West University for their supports during collection of samples and data from the wheat fields. We also thank Dan MacLean, Emilie Chanclud, and members of the Kamoun Lab for their suggestions, the staff of the Biotechnology Biological Sciences Research Council (BBSRC) National Capability in Genomics at TGAC including Christopher Wright, Helen Chapman, Harbans Marway, Tom Barker, and Jonathan Moore for assistance with the sequencing. Authorization for scientific activities #39131-3 from the Brazilian Ministry of Environment (MMA) / “Chico Mendes” Institute for Conservation of Biodiversity (ICMBIO) / System for Authorization and Information in Biodiversity (ICMBIO) for studying the evolution of *Pyricularia oryzae* from wheat, issued to VLC and PCC. This work was funded in part by the World Bank under a HEQEP CP # 2071 to MTI, a BBSRC fellowship in computational biology awarded to DGOS, the NBI Computing infrastructure for Science (CiS) group, Brazilian National Council for Scientific and Technological Development – CNPq (Pq-2, 307295/2015-0) and São Paulo Research Foundation – FAPESP (2014/25904-2, 2013/10655-4 and 2015/10453-8) research grants to VLC and PCC, and the Gatsby Charitable Foundation and BBSRC to JW and SK.

## AUTHOR CONTRIBUTIONS

MTI, JW, SK: Conceived and coordinated project, designed and performed data analyses, wrote paper. DGOS: Conceived and coordinated project, designed and performed analyses, performed experiments, revised paper and/or provided scientific insight. DC, PG: Designed and performed analyses, wrote paper. DMS, MGM, PCC, NJT: Collected, processed, and contributed samples/data, revised paper and/or provided scientific insight. AP, PB, SH, DRG: Collected, processed, and contributed samples/data, performed experiments. MMR, MUS, VBS, JLNM, ANJ, VLC, JTdAR, SR, TS, DS: Collected, processed, and contributed samples/data. NC: Performed experiments. RK, EF, DT, MHL, BAM: Revised paper and/or provided scientific insight.

